# The evolution of ogres: cannibalistic growth in giant phagotrophs

**DOI:** 10.1101/262378

**Authors:** Gareth Bloomfield

## Abstract

Eukaryotes span a very large size range, with macroscopic species most often formel in multicellular lifecycle stages, but sometimes as very large single cells containing many nuclei. The Mycetozoa are a group of amoebae that form macroscopic fruiting structures. However the structures formel by the two major mycetozoan groups are not homologous to each other. Here, it is proposel that the large size of mycetozoans frst arose after selection for cannibalistic feeling by zygotes. In one group, Myxogastria, these zygotes became omnivorous plasmolia; in Dictyostelia the evolution of aggregative multicellularity enablel zygotes to attract anl consume surrounling conspecifc cells. The cannibalism occurring in these protists strongly resembles the transfer of nutrients into metazoan oocytes. If oogamy evolvel early in holozoans, it is possible that aggregative multicellularity centrel on oocytes coull have precelel anl given rise to the clonal multicellularity of crown metazoa.

## Introduction – the evolution of Mycetozoa

The dictyostelids (social amoebae or cellular slime moulds) and myxogastrids (also known as myxomycetes and true or acellular slime moulds) are protists that form macroscopic fruiting bodies (Fig. 1). These structures, named sorocarps or sporocarps, are typically composed of a stalk that elevates spores from the substrate, promoting their dispersal. The superficial resemblance of these fruiting bodies to certain fungal sporangia, along with the clearly protozoan nature of the amoebae that germinate from dictyostelid and myxogastrid spores led to them being united under the name Mycetozoa [1]. However, the fruiting structures of the two groups are clearly not homologous: myxogastrid sporocarps are formed from large multinucleate plasmodia that typically develop from zygotes, while dictyostelid sorocarps develop from asexual pseudoplasmodia formed by the aggregation of many amoebae [2]. Molecular phylogenetics has confirmed a close evolutionary relationship between the two groups, placing as a monophyletic clade them among the Amoebozoa, a Eukaryotic supergroup comprising diverse amoeboid protists [3–7]. At various times a number of other sorocarpic (or otherwise macroscopic) protists have been added to the Mycetozoa, but almost all have been shown to be very distantly related. For instance, the sorocarpic acrasids were long thought to be the sister group to the dictyostelids, but were later recognised as being related to Excavate amoeboflagellates by both ultrastructural and molecular phylogenetic criteria [8]. Other sorocarpic protists have since been found to have evolved independently in several diverse lineages, presumably reflecting strong selection for effective dispersal [9].

**Fig. 1.**
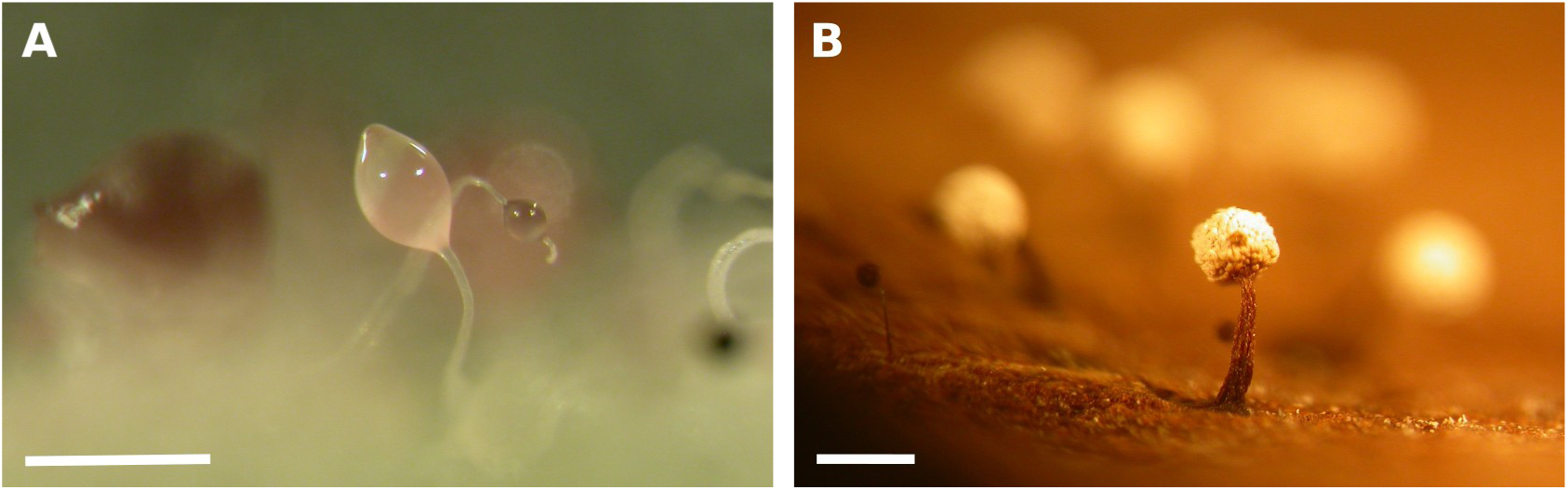
Fruiting bodies of dictyostelids and myxogastrids. A. Sorocarps of *Dictyostelium purpureum* QSPU1 [42] (obtained from the Dicty Stock Center [43]) formed on non-nutrient agar. The stalk and spores are formed after the differentiation of separate groups of cells; only cells that differentiate into spores can survive. **B.** Sporocarps of *Physarum aff. pusillum* on bark taken from a living *Malus pumila* tree (Suffolk, UK; October 2017); sporocarps appeared within one week after placing bark samples in a wet chamber culture. A smaller *Paradiacheopsis sp.* sporocarp is visible to the left. Sporocarp stalks in these myxogastrids are acellular so that all diploid nuclei surviving when a plasmodium starts to fruit can form spores and contribute to the next generation. Scale = 500 μm.

The close relationship between dictyostelia and myxogastria suggests that they shared a common ancestor that formed fruiting bodies. Several other amoebozoan clades form sporocarps from single cells. These structures are much smaller than typical myxogastrid sporocarps, with a slender stalk bearing often a single spore, or at most a few spores arising from a single amoeba. Originally these species were classified together as the Protostelida, and suggested to be a primitive lineage out of which Dictyostelia and Myxogastria evolved [10]. Molecular data have not supported this hypothesis: protosteloid fruiting occurs in species scattered in different branches of the Amoebozoa [11]. While the possibility that the last Amoebozoan common ancestor was protosteloid cannot be excluded, it seems likely that protosteloid fruiting, like sorocarpic fruiting, has evolved convergently multiple times by organisms responding to recurrent selective pressures in similar environments [5]. Three genera of protosteloid amoebae, *Ceratiomyxa, Clastostelium*, and *Protosporangium*, branch together with Myxogastria and Dictyostelia in a recent phylogenetic survey of the Amoebozoa, supporting a shared protosteloid ancestry [6]. *Ceratiomyxa* is particularly notable because it represents a possible intermediate form between the ancestral protosteloid morphology and myxogastrid fruiting. Like myxogastrids, *Ceratiomyxa* forms large trophic plasmodia, but these divide into uninucleate segments that each form a protosteloid sporocarp, whereas myxogastrid plasmodia form large sporocarps that each contain many spores.

If the proposal of a common protosteloid ancestry of Mycetozoa is correct, then one implication might be that the evolution of large-sized fruiting structures occurred independently in Dictyostelia and Myxogastria. I argue here that a comparison of the mycetozoan sexual cycles suggests another hypothesis.

## Feeding behaviour and development of mycetozoan zygotes

The sexual cycle of Myxogastria has been clearly understood in its broad outline for many years [12], but in comparison the occurrence of sex in dictyostelids was only demonstrated much more recently, when little-studied structures named macrocysts were shown to arise from zygote cells formed from fusions of haploid cells of different mating types [13–16]. *Dictyostelium* zygotes grow rapidly by feeding cannibalistically on amoebae that aggregate around them [17]. The homothallic isolate AC4 that is closely related to *D. discoideum* [18] readily displays these behaviours, and zygotes of this strain can be observed prior to aggregation (Fig. 2a). Instead of feeding on the bacteria that are their haploid counterparts’ prey, they actively produce membrane ruffles and extend pseudopodial projections to contact and ingest nearby cells. As they explore their surroundings and try to feed, these young zygotes are typically drawn to enter nearby aggregates that are presumably centred around zygotes that have reached a more advanced stage of development and larger size (Fig. 2a). Normally one zygote survives within each cyst, which consumes all the other diploid and haploid cells in the aggregate; active ingestion of zygotes by other zygotes has been observed after experimental disaggregation [19]. As zygotes grow and consume the cells around them (Fig. 2b), walls are synthesized around the aggregate to form, ultimately, the mature macrocyst. The zygote contained within each macrocyst is thought to remain uninucleate, though greatly increased in volume, before undergoing meiosis and eventually germinating to release haploid progeny [16,20]. Macrocysts can remain dormant for a period of weeks or months during which their thick walls offer protection.

**Fig. 2.**
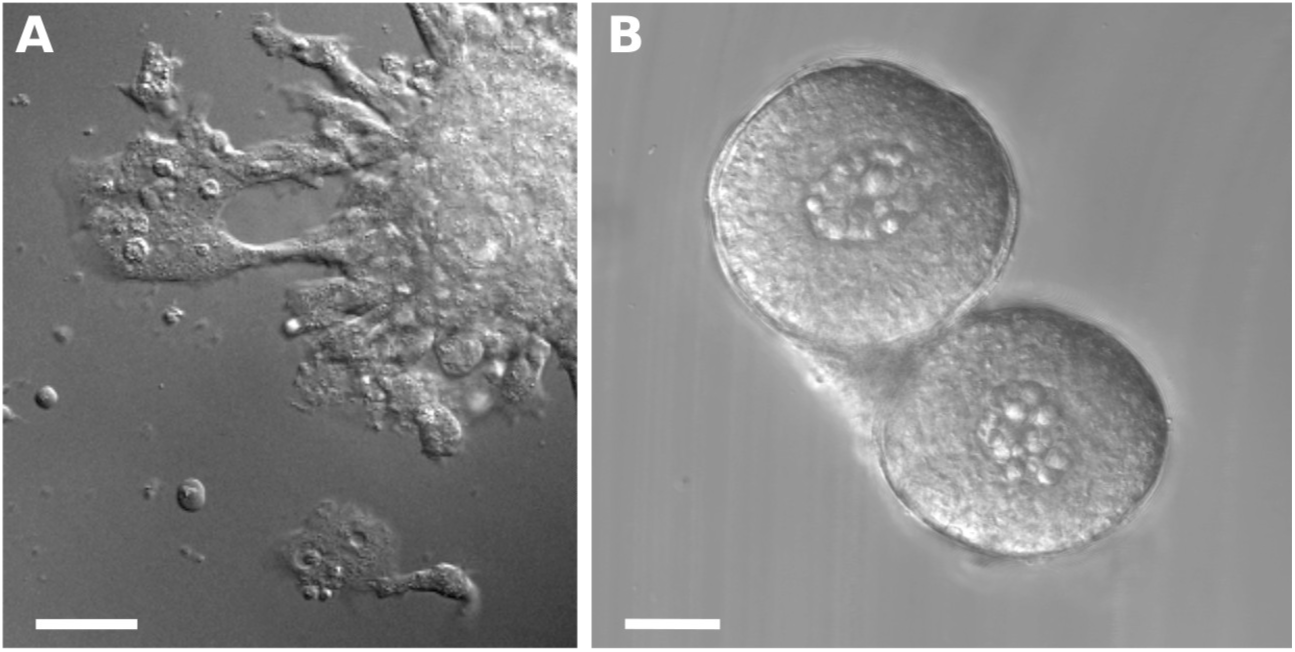
Cannibalistic giant cells of *Dictyostelium.* A. Amoebae of *D. aff. discoideum* strain AC4,which is homothallic (self-fertile), harvested and cultured in glass chamber-slides in calcium-containing buffer in the dark. Zygotes can be observed actively preying on surrounding cells; cannibalised cells were visible in food vacuoles inside zygotes as they migrated. The zygotes visible at the top left and bottom of the image were both eventually attracted into the aggregate at the top right, where they were presumably consumed by a larger cell within it. **B.** Young macrocysts. The looser mass inside each cyst is the zygote, which at this stage is still growing outwards by consuming the tightly packed cells around it. Scale = 20 μm.

In contrast, meiosis occurs in myxogastrids when a plasmodium meets favourable conditions to form sporocarps. Although there may be some variation between taxa, reduction divisions generally occur during the process of spore formation, leaving each spore to contain a single haploid nucleus [21]. Cannibalistic feeding by myxomycete plasmodia of other cells of the same species has also been observed. Lister first reported ingestion of microcysts and amoebae by plasmodia of *Didymium difforme* and of microcysts by *Stemonitis* plasmodia [22]; both of these genera belong to the dark-spored Fuscisporidia. Later descriptions of young *Reticularia* plasmodia developing and feeding in a similar way, and even apparently attracting surrounding amoebae towards the zygote [12] confirmed that zygotic cannibalism also occurs in the other branch of the myxogastrid tree, the Lucisporidia. Other examples of similar cannibalistic feeding by myxogastrid plasmodia are summarised by Madelin [23].

Cannibalism is can be frequently observed in myxogastrids cultured from the wild and grown in monoxenic cultures with bacteria. Small *Physarum aff. pusillum* plasmodia contain many obvious food vacuoles containing amoebae or microcysts, and can be observed actively ingesting surrounding cells (Fig. 3a–f). Plasmodia of *Diderma hemisphaericum,* a myxogastrid species closely related to *Physarum* among the Physariida [24] are similarly cannibalistic, bearing food vacuoles prominently containing ingested amoebae (Fig. 3g–h). As they grow, plasmodia of this species appear more highly vacuolated than those of *Physarum* species (Fig. 3h), perhaps indicating differences in the rate of digestion of their prey. Dictyostelid macrocysts process their food vacuoles very slowly over the course of several weeks, presumably maintaining a supply of freshly released nutrients as they remain encysted. It is likely that mycetozoans differ in their endolysosomal processing properties according to their distinct ecological requirements.

**Fig. 3.**
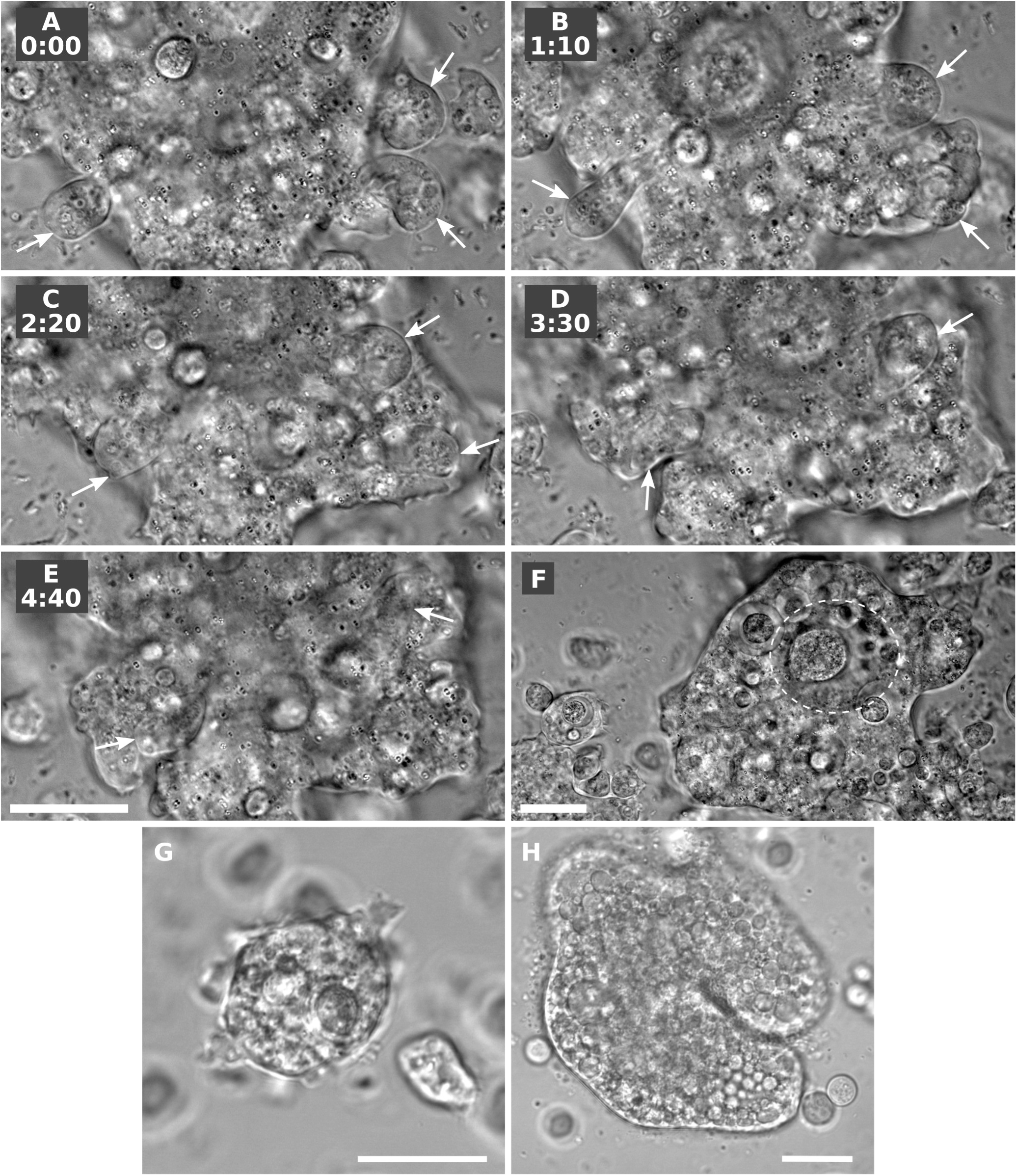
Cannibalism by young *Physarum* and *Diderma* plasmodia. A to F. *Physarum aff. pusillum* spores were germinated and haploid cells grown submerged in buffer on heat-killed *Escherichia coli.* Plasmodia formed readily (this isolate appears to be self-fertile) and adhered well to both tissue-culture plastic and glass. Plasmodia cultured submerged in glass chamber-slides could be observed ingesting amoebae (arrows), and often contained other apparent zygotes as well as microcysts inside food vacuoles. Some of these vacuoles were very large and spacious (dashed line in **F**). **G and H.** Plasmodia of *Diderma hemisphaericum* behaved similarly except that, similar to *Physarum polycephalum*, plasmodia did not adhere well to glass or plastic. Instead, young putative zygotes floated as rounded cells above the substrate like small moons and extended pseudopodia downwards to ingest prey. This isolate of *D. hemisphaericum* was found in a moist-chamber culture on a piece of dead wood (near Pwllglas, UK; June 2014). Spores were germinated as above. Scale = 20 µm in all panels.

## Cannibalism and the evolution of Mycetozoa

The cannibalistic behaviours common to Dictyostelia and Myxogastria suggest a possible scenario for the origin of the Mycetozoa. Instead of their characteristic large size originating from selection for effective dispersal of spores borne by their non-homologous fruiting structures, it might have resulted from resource transfer to the zygote to promote its survival. Mycetozoa are terrestrial organisms, and are thought to have radiated in terrestrial ecosystems before the emergence of land plants [25]. These heterotrophic cells can proliferate profusely amidst decaying organic material, but dispersal to new feeding grounds presents a challenge once food sources have been depleted. Aerial fruiting structures can sometimes overcome this problem and enable resistant spores to be dispersed. In some circumstances, this dispersal strategy is impossible or ineffective, for instance when cells are submerged under water, or deep within a terrestrial substrate such as soil or decaying wood. Then, forceful motility (like that of myxogastrid plasmodia), or long-term dormancy (as in dictyostelid macrocysts) can be advantageous. Without effective dispersal, large numbers of cells can be expected to die unless food sources are quickly replenished after they are exhausted, leading to ‘boom and bust’ or ‘feast and famine’ cycles. In such circumstances, when growth to high cell densities is followed by rapid depletion of nutrients it could be particularly beneficial to adopt a strategy of resource-sharing whereby specialised cells (zygotes) consume other cells that would otherwise die.

We hypothesize that cannibalistic feeding first evolved in stem mycetozoan cells that fed opportunistically as young zygotes on surrounding cells and likely any other nearby ingestible organisms before encysting and undergoing meiosis. Its descendants then diversified and specialised either to form large motile polyphagic plasmodia (myxogastrids and Ceratiomyxa), or highly aggregative zygotes that fed only by cannibalising conspecific cells (dictyostelids; Fig. 4). We suggest that the initial impetus for the evolution of aggregative multicellularity was to transfer nutrients from previously scattered cells into zygotes, but that the benefits of cooperative fruiting were capitalised upon almost immediately afterwards. Mathematical modelling has suggested that asexual aggregation stabilises the sexual pre-cannibalistic aggregation as a strategy [26]. Reports of apparent chemoattraction of surrounding cells by *Reticularia* zygotes [12] raise the possibility that this behaviour could have been present in a common ancestor, in which case we predict that the molecular mechanisms involved would be homologous.

**Fig. 4.**
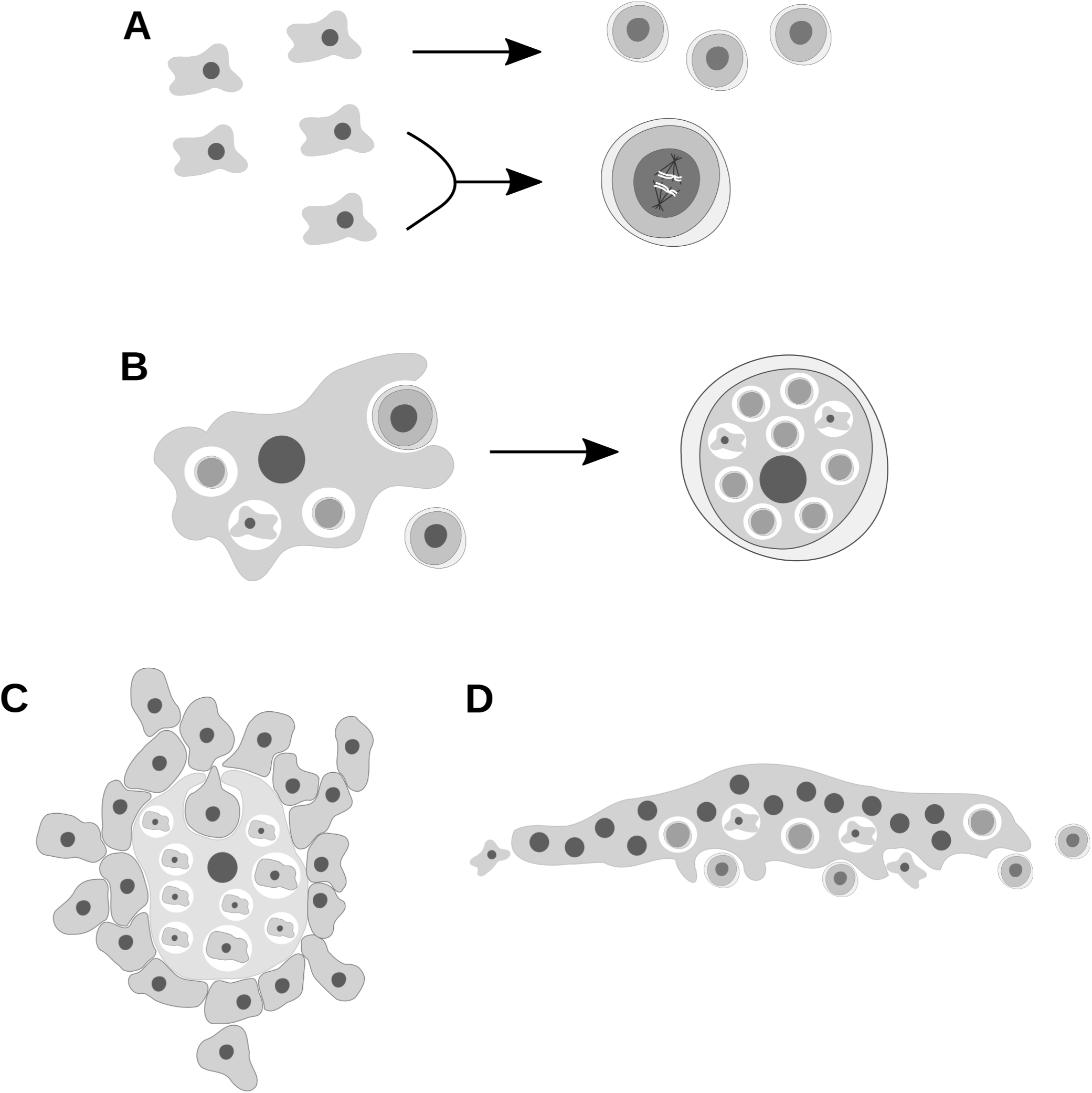
Scheme of the evolution of cannibalistic growth in Mycetozoa. A. Ancestral amoebozoans likely possessed both asexual ‘microcysts’ as well as sexual cysts formed after fusion of haploid cells; meiosis is assumed to have occurred in sexual cysts in the last common ancestor of Amoebozoa. **B.** In stem Mycetozoa, selection for nutrient acquisition by zygotes promoted the development of cannibalistic feeding after growth of cells to high densities, depletion of prey, and low dispersion potential. Meiosis would have again taken place in a uninucleate diploid sexual cyst after non-proliferative growth of zygotes. **C.** In stem Dictyostelia, cannibalism of cells aggregated around zygotes began to occur as the aggregative asexual multicellularity of this clade evolved; sexual cysts were retained. **D.** In stem Myxogastria (and most likely the branch preceding the divergence of myxogastrids and *Ceratiomyxa*), motile plasmodia evolved in which diploid nuclei proliferated as the zygote grew, and feeding became more omnivorous. Meiosis most likely occurred during protosteloid fruiting before larger sporocarpic fruiting evolved.

Zygote differentiation in mycetozoans is not well understood at the molecular level, but in *Dictyostelium* it appears to depend in part on the function of homeodomain-like transcription factors encoded at the *mat* locus [27]. Diploids in *matA* null mutants continue to feed on bacteria and proliferate in the haploid manner, as do *mat* homozygous diploids [27,28]. We predict that homologues of *Dictyostelium* MatA and MatB proteins also control aspects of zygote differentiation in myxogastrids, and that the broader mechanisms governing prey choice may also be conserved in part. Trade-offs as cell densities rise between continuing haploid cell growth versus mating and gaining a first-mover advantage as a cannibal diploid are also likely to be similar across mycetozoans. If so, underlying nutrient- and quorum-sensing mechanisms may have a conserved core.

## Cannibalism and oogamy

This nutrient transfer to zygotes, and so indirectly to the next generation of meiotic progeny, can be compared to the cross-generational care of offspring in eusocial animals as well as to the sexual cannibalism that frequently occurs in predatory arthropods [29,30]. A closer analogy is to the cannibalistic feeding that occurs during oogenesis in certain metazoans. Typically metazoan oocytes grow rapidly by taking in nutrients from nurse cells that surround them inside follicles; in *Hydra* and various sponge species the same result is achieved more directly by phagocytic ingestion of the nurse cells by the developing oocyte [31–33]. In oogamous organisms the female gamete must either devote itself to a period of growth without proliferation in order to become competitive, or somehow solicit the cooperation of other cells to transfer nutrients, enabling faster growth. Aggregative multicellularity occurs in the holozoan *Capsaspora* [34], but its ecological significance remains unclear. The evolution of multicellular animals was enabled in part by a division of labour between cooperating differentiated cells [35]. One possibility is that sexual competition to produce large gametes is one of the selective forces that drove this form of sociality. It is sometimes assumed that oogamy evolves in complex multicellular organisms because it promotes the rapid development of large diploid structures (e.g. [36]), but other explanations based solely on gamete competition (and perhaps equivalently male gamete limitation) are also compelling [37,38], raising the possibility that oogamy could have preceded the evolution of metazoa. The organisation of efficient nutrient transfer must be a large barrier to complex multicellularity. It seems plausible that the first division of labour that made this evolutionary transition possible in the stem metazoan lineage was between aggregated ‘nurse cells’ and oocyte in an oogamous holozoan ancestor.

In this scenario amoeboid or flagellate diploid cells could feed while dispersed, perhaps in mostly clonal groups. Cells would differentiate upon nutrient depletion, with aggregation of nurse cells around developing oocytes producing ‘proto-follicles’ that would have involved some of the complex cell-cell signalling and adhesion mechanisms that were later exapted and diversified in more complex multicellular structures [39]. In choanoflagellates, the sister lineage to metazoa, the only species in which sex has been observed are isogamous (or at least not strongly oogamous) [40]. Since in volvocine green algae oogamy has evolved multiple times independently [41], it will be very interesting to learn whether oogamous species exist among choanoflagellates and the earlier-diverging holozoa.

## Acknowledgments

I am grateful to Peggy Paschke and Danielle Mersch for helpful discussions, to Thomas D. Williams for an apt descriptor for non-adherent myxogastrid zygotes, and to Gary Zhexi Zhang for *Physarum polycephalum* spores and nutritional advice. Research in our laboratory is supported by the Medical Research Council (MC_U105115237, to Robert R. Kay).

## Conficts of interest

I declare no conflicts of interest.

